# Gut microbes of the cecum versus the colon drive more severe lethality and multi-organ damage

**DOI:** 10.1101/2024.02.26.582076

**Authors:** Kejia Xu, Juan Tan, Dongyang Lin, Yimin Chu, Luting Zhou, Junjie Zhang, Yinzhong Lu

## Abstract

An intestinal perforation or puncture leads to a high risk of sepsis-associated morbidity. A perforation initiates the transfer of the intestinal contents (ICs) to the peritoneal cavity, leading to abdominal infections and varying with different prognoses. However, the mechanisms associated with different perforations in the cecum and colon remain unknown. We sought to examine how different gut flora contribute to prognoses in different intestinal perforation sites. We compared the microbiome of the ICs in the cecum and colon in a fecal-induced peritonitis mouse model. The results showed that cecum ICs developed more severe sepsis than colon ICs, including a shorter median survival time, increased biochemical indicators, more pathological changes in multiple organs and overwhelmed systematic inflammation. Moreover, our results demonstrated that cecum ICs hold more bacterial burden in unit weight than colon ICs, and the microbial communities differed between the ICs from the cecum and colon. A more detailed comparison of the two microbiome groups showed that the abundance of potentially pathogenic bacteria increased in the cecum ICs. Our data suggest that the sepsis severity developed by perforation was associated with bacterial burden and increased abundance of potentially pathogenic bacteria in the cecum. Our findings first compared the differences in the lethality associated with the ICs of the cecum and colon, which pointed out that the site of perforation could help providers predict the severity of sepsis.

## 1 Introduction

Abdominal infections commonly occur after perforation of the lower digestive tract. Lower digestive tract perforation, mainly including cecal perforation and colonic perforation, is a prevalent but acute disease that threatens life. The leading cause is intestinal contents (ICs) leakage to the sterile peritoneal cavity, thus contributing to harmful or lethal infection, severe multi-organ damage, and even death [1, 2]. Such perforations can increase sepsis-related mortality and morbidity [3–5]. Once a perforation occurs, emergency surgery and antibiotic treatment are required, but its prognosis varies with the patient’s overall health status, diagnosis, and surgery duration [3, 6]. Some retrospective studies showed that cecal and colon perforations have different prognoses [7–10], which implies a different mechanism of lethality. However, clinical retrospective pilot studies have drawn inconsistent conclusions for which different outcomes vary in different sites of the lower digestive tract [11, 12]. Therefore, these clinical issues are still being debated and need further clarification.

Gut microbes vary and can be drivers of many diseases in previous studies [13–17], which can predict patient susceptibility to disease and provide microbiota-targeted tools for disease treatment [18]. For example, the fecal microbiome signature exhibits high specificity for colo<u>n</u> cancer [19], non-alcoholic fatty liver disease development (NAFLD) [20], and Parkinson’s disease [21]. Interestingly, these microbiome signatures produced strategies for treating Parkinson’s disease [21], ulcerative colitis [22], and intestinal inflammation [23]. In colonic perforation-associated sepsis treatment, empiric antibiotics are routinely prescribed, but their efficacy varies because no specific antibiotics or spectrum of antibiotics to treat sepsis are available [1, 17] due to uncontrolled bacterial infection. One possible reason is that the patient misses an optimal treatment window when the physicians are forced to modify the therapy plan to tackle the unsatisfactory results of empiric antibiotic therapy [24], which implies it is wise to target the driving bacteria. It reports that gut microbe compositions vary in their different fragments of the low digestive tract in humans and animals [25, 26], and several lethal or infectious bacterial strains were previously reported in gut microbiota [2, 27]. Therefore, it can be reasonably considered that sepsis caused by colonic perforation needs further analysis, and whether it is driven by different gut microbiota remains to be explored.

Intra-abdominal sepsis hospital admissions took up 20% of patients with sepsis [28], which could be mimicked with polymicrobial rodent models for basic research [29]. The polymicrobial models of sepsis are typically applied as they involve either cecal ligation and puncture (CLP) or fecal-induced peritonitis (FIP) with intra-peritoneal injection. Nevertheless, in the CLP model with an open abdominal setup, the cecum is ligated and punctured with a needle to allow ICs to leak into the sterile peritoneal cavity [30]. Thus, the CLP model introduces the risk of infection caused by surgery that would add additional complicated factors to the outcome. Instead, FIP provides an alternative model for mimicking the polymicrobial sepsis model without surgery [29]. Therefore, the FIP model might be reasonably used to evaluate the severity of the on-site ICs in sepsis development.

In the study, we sought to examine whether gut microbiome from different perforation sites in the intestine would cause variations in sepsis development. We used a mature FIP sepsis model for the first time to evaluate the effects of ICs from different sites (cecum or colon) on the outcomes by monitoring their survival, blood biochemical indicators, systemic cytokines, lung inflammation, and histological (liver, lung and kidney) alterations of the model mice. In addition, we compared the microbiomes of the cecum and colon and linked their bacterial number and microbial community structure to the severity of perforation sites.

## 2. Results

### 2.1 Different survival time between the mice receiving ICs from the cecum and colon

As shown in **Figure 1A**, we monitored the survival time following the injections with IC solutions from different intestinal sites. The Kaplan–Meier curve showed different survival times between mice receiving IC aliquots from the cecum and colon. The ICs from the cecum group showed a much shorter median survival time than the colon group (22.57 *vs.* 25.20 h; p < 0.0001, **Figure 1B**), while no deaths were observed in saline-injected mice. These results demonstrate that cecal ICs exerted more lethal effects than those from the colon.

**Figure 1.**
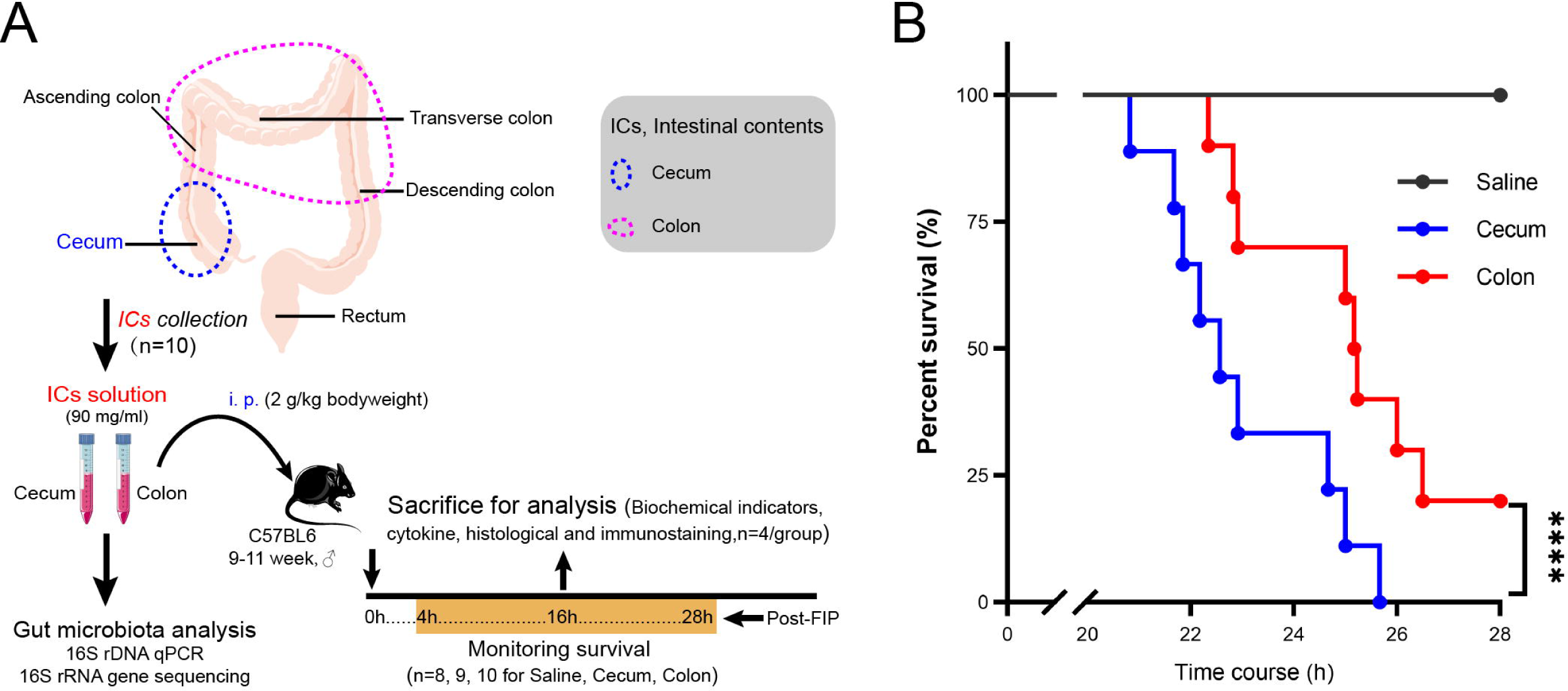
Study design and survival rates monitering. (A) Study design including treatment and sampling times. (B) Survival rates during the fecal-induced peritonitis (FIP)-induced sepsis. Saline, cecum, and colon show the mice injected intraperitoneally with their respective saline or IC aliquots. ****P < 0.0001, indicating a significant difference between the cecum (n=9) and colon (n=10) based on the Mantel–Cox test.

### 2.2 ICs from different intestinal sites induce different blood biochemical indicators

The blood biochemical indicators exhibit the degree of organ injuries in rodents and humans. Thus, biochemical blood indicators were detected and demonstrated various degrees of injuries among the cecum-FIP and colon-FIP groups. We noted that biochemical indicators of impaired liver function, alanine transaminase (ALT), tended to increase in the cecum group compared to the control or colon group but without a noticeable difference (p=0.24 and 0.34, respectively, **Figure 2A**). The aspartate transaminase (AST) of the cecum-FIP group increased when compared to the control group (p<0.05, **Figure 2B**) and tended to be higher than the colon group (p=0.10, **Figure 2B**). In addition, an indicator of tissue damage lactate dehydrogenase (LDH, p<0.05; **Figure 2C**), and indicators of kidney dysfunction creatinine (p<0.05; **Figure 2D**), creatinine kinase (p<0.05; **Figure 2E**), and blood urea nitrogen (BUN, p=0.072, **Figure 2F**), were significantly higher in the Cecum-FIP group than the colon-FIP group. Of note, blood LDH (p<0.05), BUN (p=0.056), Creatinine (p=0.11), and CK (p=0.12) in the cecum-FIP group but not the colon-FIP group tend to increase than the control groups. The blood biochemical indicators demonstrate a more severe organ injury in cecum-FIP mice than in the colon-FIP group.

**Figure 2.**
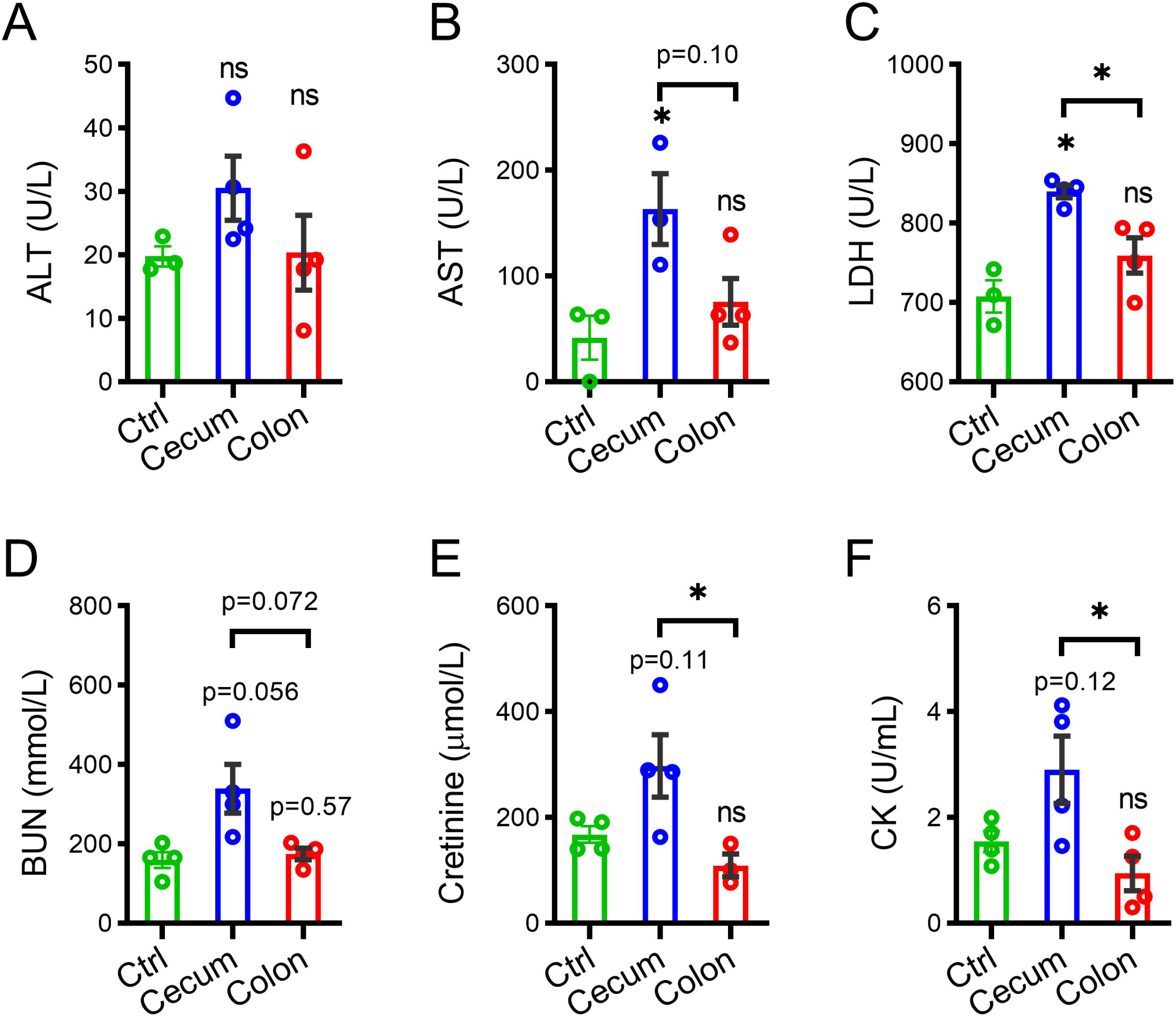
Blood biochemical indicators altered during the FIP-induced sepsis. **(A-F)** Mice were intraperitoneally injected with saline or ICs from fecal slurry (2 mg/kg) or colon slurry (2 mg/kg), and 16 hours later, plasma was analyzed for amount of (A) alanine aminotransferase (ALT), (**B**) aspartate aminotransferase (AST), (**C**) Lactate dehydrogenase (LDH), (**D**) Blood urea nitrogen (BUN), (**E**) creatinine, and (**F**) creatine kinase (CK) (each dot represents one mouse, n = 3 to 4 per group); Data represent as means ± SEM with p-value or not, \**P* < 0.05, ns, no significant; one-way ANOVA with post-t-test.

### 2.3 ICs from the cecum induce high levels of blood cytokine productions

To monitor the systemic immune response to the cecum-FIP or colon-FIP challenge at 16h, the multiplex cytokines panel was employed to show that pro-inflammatory cytokines such as TNF-α, IL-1α, IL-1β, and IL-6 in the cecum-FIP group have a higher level than the colon-FIP group (**Figure 3**). Furthermore, other plasma cytokines also elicited similar higher levels in the Cecum-FIP group than in the colon group. These data indicate an extreme immune response to the FIP developed by the cecum compared to the colon group.

**Figure 3.**
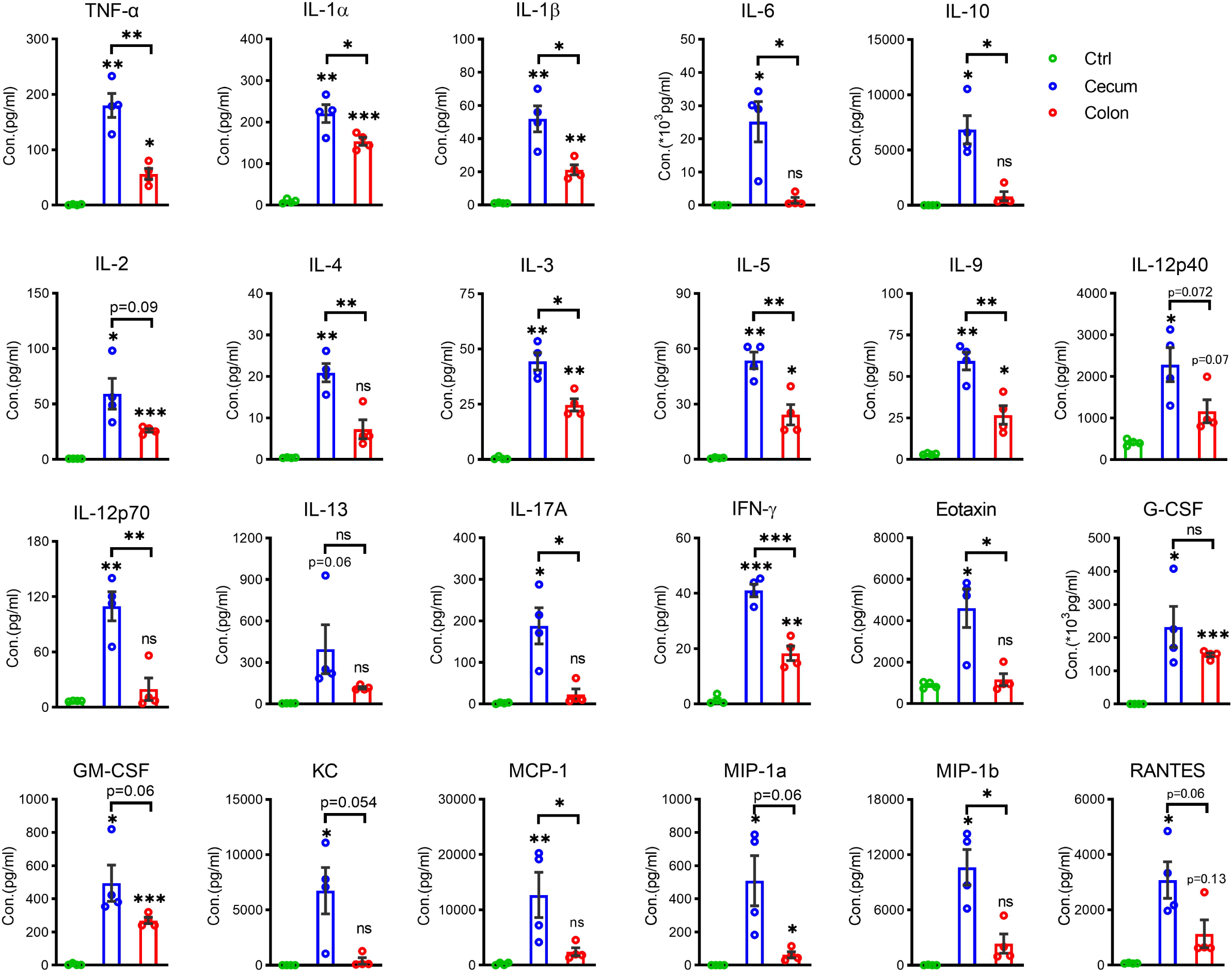
Blood plasma cytokines and chemokines increased during FIP-induced sepsis. Mice were intraperitoneally injected with saline or ICs from fecal slurry (2 mg/kg) or colon slurry (2 mg/kg), and 16 hours later, plasma was analyzed for the amount of cytokine or chemokine levels (each dot represents one mouse, n = 3 to 4 per group). Data are shown as means ± SEM. p value are shown or not, *p < 0.05, **p < 0.01, ***p < 0.001, ns, no significant; one-way ANOVA with post-t-test.

### 2.4 ICs from the cecum induce severe liver, lung and kidney pathological alterations

To evaluate the organ injuries of mice receiving ICs injections, pathological examinations of the multiple organs (liver, lung, and kidney) were found to demonstrate various degrees of pathological changes at 16 h after mice received injections with either saline or ICs from the cecum or colon (**Figure 4A**). In liver tissues, we observed that parenchymal cells demonstrated turbidly (or edema) and were swollen, and the sinus cord structure was unclear (the liver sinuses became narrowed, and a thickened disorganized liver plate was detected). Nevertheless, the cecum group had degenerated and showed balloon-like changes—accounting for ∼60% of the area of the liver sections—was scattered with individual lymphocytes and the focal point necrosis of hepatocytes could be observed (**Figure 4A**, upper panel). The Knodell scoring on the degenerative and necrotic cells and Ishak scoring on the fibrosis showed that the cecum-FIP mice tended to receive slightly higher marks than the colon-FIP mice (p=0.38 and 0.31, **Figure 4B**). In mice with ICs, the lungs showed broken focal alveolar septa, fused alveolar cavity, slightly widened focal alveolar septum, and alveolar epithelium hyperplasia compared to the control group. However, the cecum-FIP group showed more lymphocyte and neutrophil infiltration in the small focal pulmonary interstitium than in the colon-FIP group (**Figure 4A**, middle panel) and tended to receive a higher clinical score (p=0.09, **Figure 4C**). In the kidney tissues, the epithelial cells of renal tubules in small foci were loose and swollen, and tiny vacuoles with foams could be observed in the cytoplasm of both groups **(Figure 4A**, bottom panel); the severity was more intense in the cecum-FIP group when compared to the control group or the colon group (p<0.01 and p=0.34, respectively, **Figure 4D**). In addition, the *NGAL* mRNA, an early marker gene for acute kidney injury, is upregulated in both the cecum- and colon-FIP mice (**Figure 4E**), while its expression in the cecum-FIP group tends to be higher than in the colon-FIP group (p=0.32). These descriptions for the hematoxylin and eosin staining and further clinical scoring (**Figures 4B-D**) demonstrated severe pathological alterations of several tissues in the cecum-FIP mice compared to the colon-FIP group.

**Figure 4.**
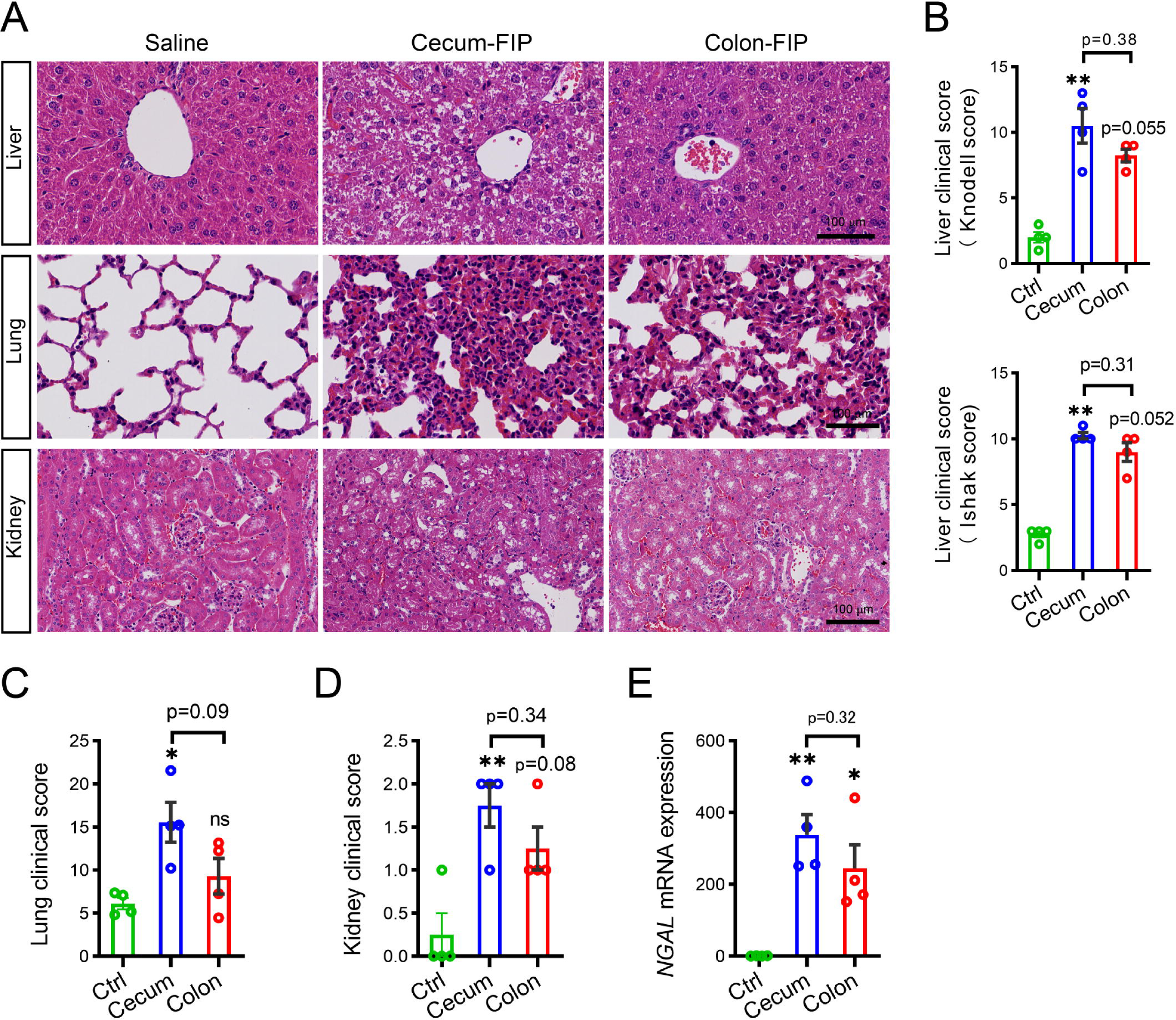
Histology altered in the cecum FIP mice. **(A)** Representative hematoxylin and eosin (H&E) staining and clinical score data for the liver, lung and kidney from the mice peritoneally injected with saline or intestinal contents (ICs) of the cecum or colon (2g/kg body weight) at 16 h post-injection. Scale bar, 100 μm; (**B-D**) The graph shows clinical scores of the liver (**B**), lung (**C**) and kidney (**D**), respectively. (**E**) *Ngal* mRNA expression measured by qRT-PCR. Data are shown as means ± SEM (n=4, each dot represents one mouse). *p<0.05, **p<0.01, and indicated p-value are shown as determined by the Kruskal-Wallis test on the clinical score and one-way ANOVA with post-t-test on the mRNA expression.

### 2.5 ICs from the cecum induce severe lung inflammation

Several indicators were stained for lung specimens to examine the lung injuries of mice receiving ICs injections. The Gr-1 (also called Ly6G) staining showed more neutrophil infiltration in the lung tissue of the cecum-FIP group than in the colon-FIP group (**Figures 5A** and **B**). Moreover, the TNF-α is more pronounced in the lung tissue of the cecum-FIP mice than in the colon-FIP mice (**Figures 5A and C**). However, the ZO-1 expression showed unobserved alterations in the cecum-FIP group compared to the colon-FIP group (**Figures 5A and D**). These data suggest that the cecum-FIP mice developed more severe lung inflammation than the colon-FIP mice, further corroborating the pathological examinations on the lung tissues (**Figures 4A, middle panel, and 4D**).

**Figure 5.**
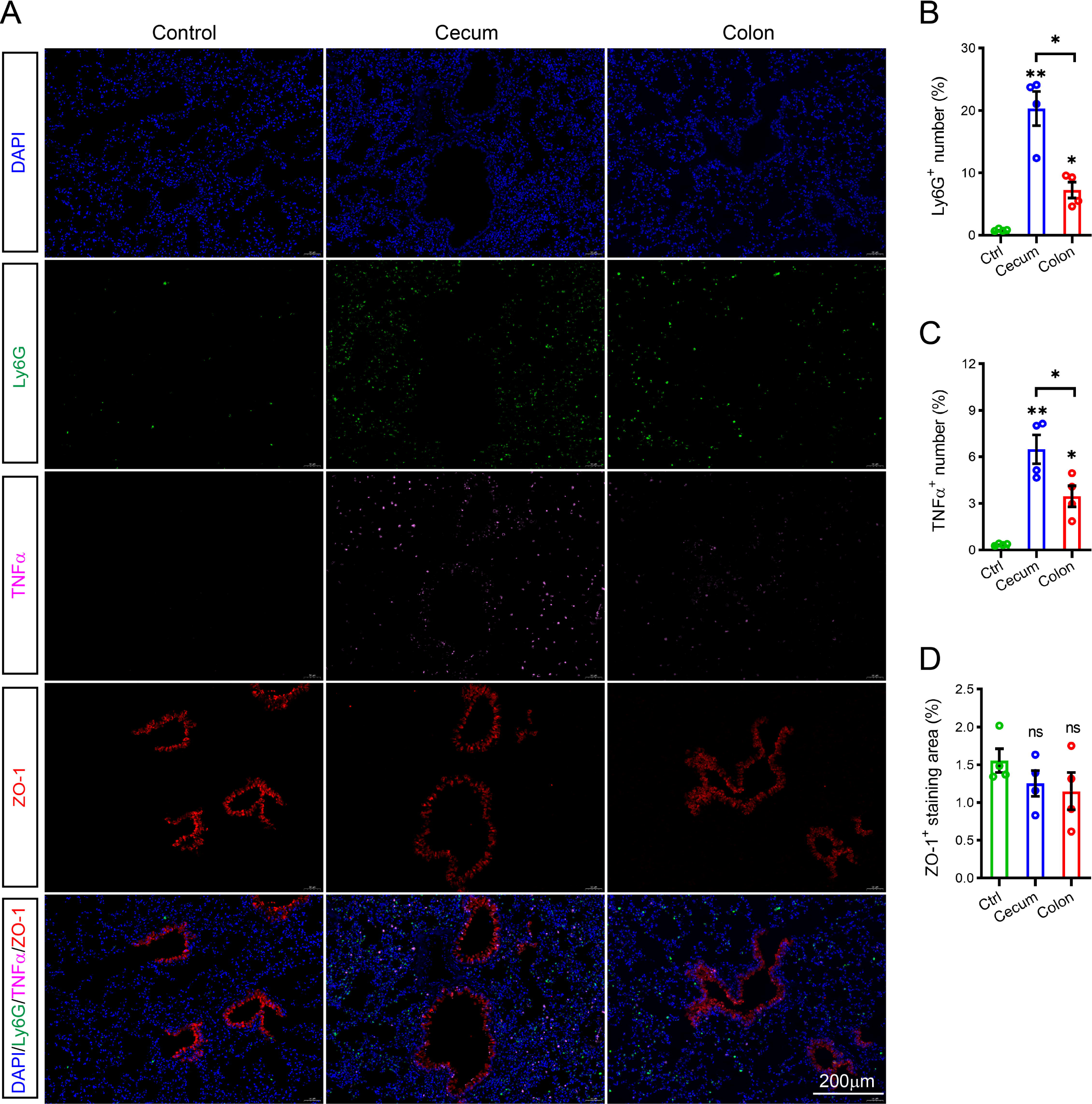
ICs of the cecum induce lung inflammation.) **(A-D** Representative image and average data for Ly6G, TNFα and ZO-1 in the lung tissues of mice peritoneally injected with saline or ICs of the cecum or colon at a dose of 2g/kg bodyweight at 16 h post-injection. Blue, DAPI; Red, ZO-1; Pink, TNFα; Green, Ly6G. Scale bar, 200 μm. The graph shows quantification data of (**A**) as the percentage of Ly6G-positive (**B**), TNFα-positive number cells (**C**) and ZO-1 staining area (**D**), respectively. Data are shown as means ± SEM (n=4, each dot represents one mouse). *p<0.05, **p<0.01, ns, no significant, one-way ANOVA with post-t-test.

### 2.6 The gut microbiome varied in different ICs

To associate the severity of sepsis with the gut microbiota, we compared the bacterial burden in the ICs by the 16S rDNA qPCR, and the quantification results demonstrated that the cecum ICs contained 3 times bacterial copy per unit weight than in the colon ICs group (**Figure 6A**). Moreover, we used 16S rRNA gene sequencing to evaluate the gut microbiota from either cecal or colonic sections. Overall, 1,286,949 effective 16S sequencing tags were obtained from the ICs of 10 young mice with an average of 64347.45 tags per sample (ranging from 56755 to 67929). After taxonomic identification, the validated tags were assigned to 98,443 operational taxonomic units (OTUs) for further analysis (**Supplementary Table S1**). According to the rarefaction (**Supplementary Figure S1A**) and species accumulation curves (**Supplementary Figure S1B**), the current sequencing and samples were sufficient for taxa identification. Furthermore, the flower plot showed that all ICs contained a core of 843 OTUs, and most could discriminate the ICs of the cecum from the colon (**Figure 6B**). Nevertheless, the rank abundance distribution curve showed no noticeable changes in the richness and evenness of the bacteria in both groups (**Supplementary Figure S1C**).

**Figure 6.**
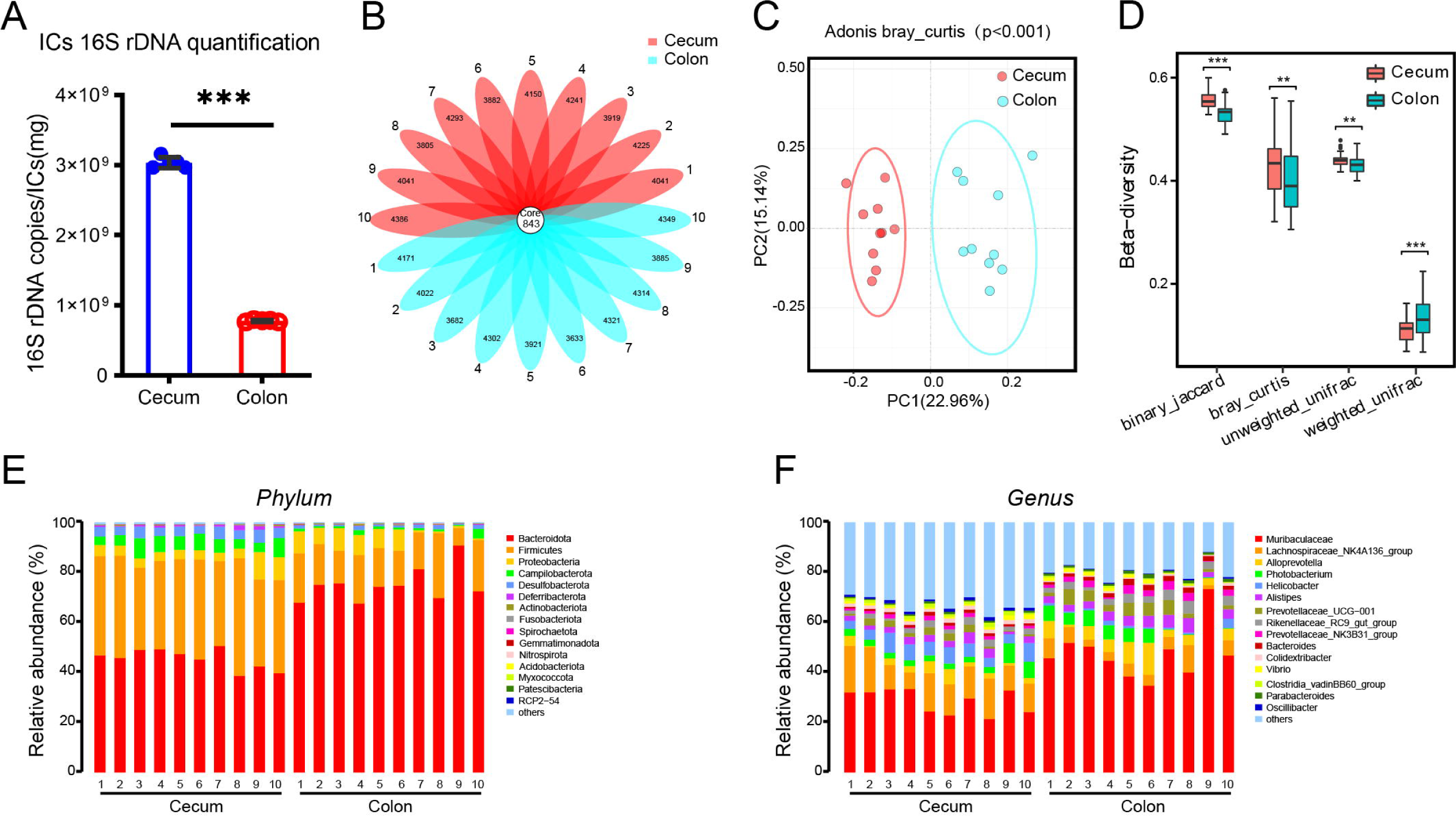
Microbiome community difference in the cecum and colon. **(A)** qPCR quantification of bacterial 16S rDNA in ICs of donor mice. Data presented as means ±SD (n = 5). (**B**) The flower plot of the operational taxonomic units (OTUs) identified in the ICs of the cecum and colon. (**C**) The principle coordinate analysis (PCoA) ordination of the Bray–Curtis distances between the cecum and colon based on Adonis bray_curtis (P < 0.001). (**D**, **E**) Relative abundance of bacterial phylum and genus in ICs from cecum and colon.

Next, we compared the microbial difference between ICs from the cecum and colon. The alpha diversity indexes, including Shannon, Simpson, Chao1, and PD tree indices in the cecal ICs, were not significantly altered compared to the colon group (**Supplementary Figure S1C–F**), suggesting similar richness and alpha diversity. However, a difference in beta diversity between the two groups was observed based on the PCoA analysis of the Bray–Curtis distance (**Figure 6C**), which was significant according to the Adonis analysis (p = 0.001). Additionally, the binary_jaccard, bray_curtis, and unweighted_unifrac distance (**Figure 6D, Supplementary Table S2**) significantly increased in the cecum group. In addition, our data showed a considerable difference in the relative abundance of each bacterium at the phylum and genus levels across samples in the cecum and colon (**Figures 6E, F, and Supplementary Figure S2**).

### 2.7 Potentially pathogenic gut microbes differ between the cecum and colon

To investigate the alterations associated with lethality or survival time, we conducted a linear discriminant analysis effect size (LEfSe) analysis. The main differences were found to be the increase of *Firmicutes* (class *Clostridia*, order *Oscillospirales* and *Lachnospirales*, family *Oscillospiraceae*), *Desulfobacterota* (Class *Desulfovibrionia*, *order Desulfovibrionales*, *family Desulfovibrionaceae*), *and Campilobacterota (class Campylobacteria*, *order Campylobacterales*, *family Helicobacteraceae*), and a reduction in the abundance of *Bacteroidota* (class *Bacteroidia*, order *Bacteroidales, family Muribaculaceae and Prevotellaceae* in the ICs of the cecum (**Figure 7A**). In addition, some differences were observed at a lower taxonomical level. For example, *Oscillospiraceae* (genus *Lachnospiraceae_NK4A136_group*) and *Helicobacteraceae* (genus *Helicobacter*) increased in the cecum. In contrast, the colon exhibited a gain of *Muribaculaceae* and *Alloprevotella* at the genus level and *Muribaculaceae* and *Prevotellaceae* at the family level **(Figure 7B, Supplementary Figure S2)**. Furthermore, our data showed that several pathogenic bacteria were highly enriched in the cecum group, while the beneficial bacteria decreased in the cecum compared to the colon group (**Figures 7C-I and Supplementary Figure S2**). Those data indicate that the gut microbiome community structure in the cecum differs from the colon.

**Figure 7.**
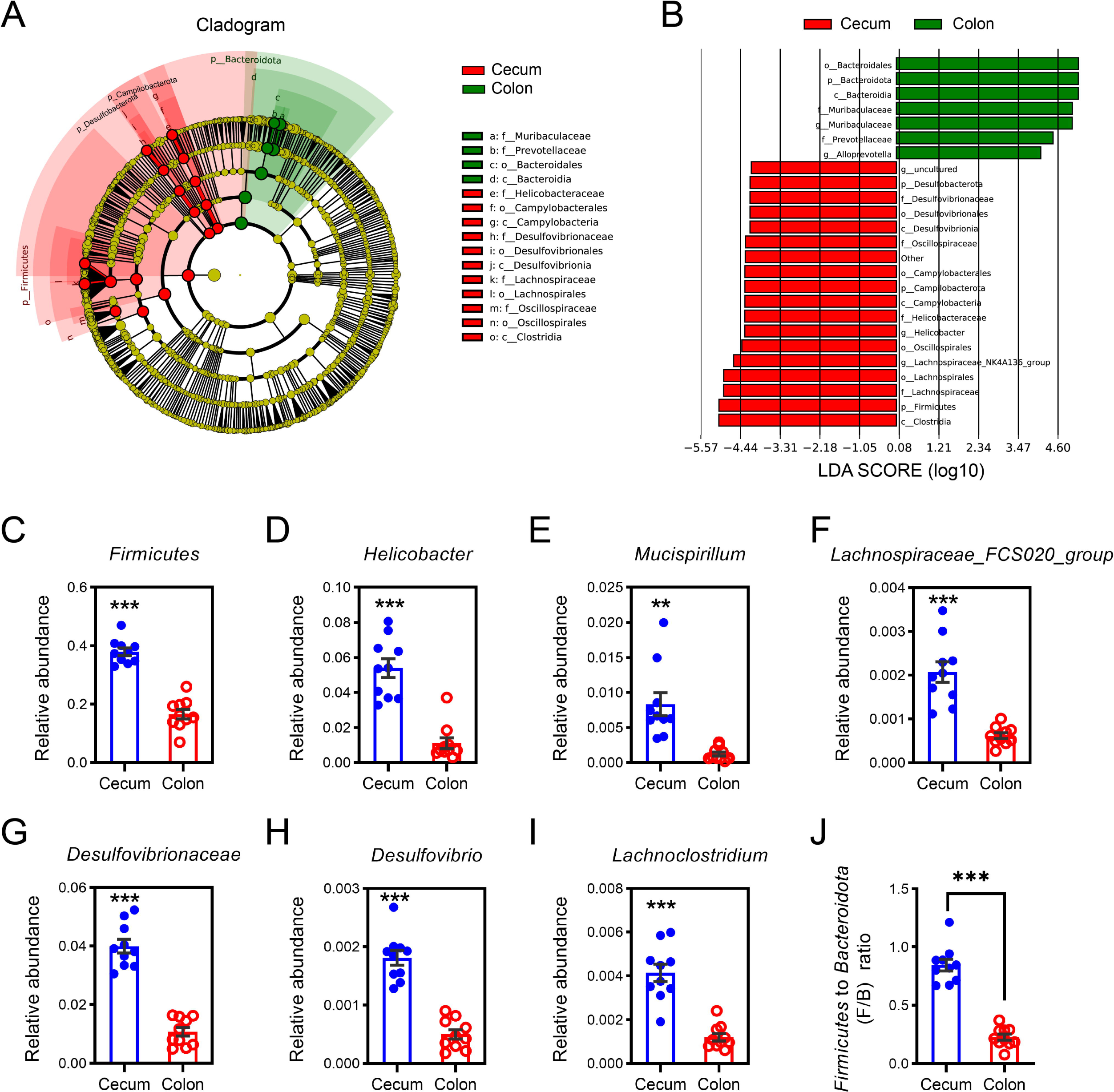
Linear discriminant analysis (LDA) Effect size analysis and the relative abundance of potentially pathogenic bacteria enriched in the ICs of the cecum. (**A**) Cladogram of the linear discriminant analysis of the effect size (LEfSe) of the microbiome from the cecum and colon ICs. Red and green circles represent the differences between the most abundant microbiome class. The diameter of each circle was proportional to the relative abundance of the taxon. (**B**) Histogram of the LDA scores for different abundant genera in the cecum and colon ICs. Red, enriched in Cecum ICs; Green, enriched in Colon ICs. **(C-I)** The relative abundance of bacteria, including *Firmicutes* (**C**), *Helicobacter* (**D**), *Mucispirillum* (**E**), *and Lachnospiraceae_FCS020_group* (**F**), the family *Desulfovibrionaceae* (**G**)*, Desulfovibrio* (**H**), and *Lachnoclostridium* (**I**). (**J**) The *Firmicutes* to *Bacteroidota* (F/B) ratio in the cecum group was higher than in the colon group.

## 3. Discussion

Intestinal perforation or puncture presents a high risk for patient morbidity, mortality, and sepsis severity. The cecum and colon were major perforation sites, especially the cecal perforation, which was more common in patients [31, 32]. Nevertheless, the ICs of different lower digestive tract fragments were variably abundant in the microbiome, food debris, metabolites, and digestive secretions. In this study, we compared the microbiome of the cecal and colon ICs and their lethality in an FIP mouse model. The results showed that the ICs of the cecum developed more severe sepsis than the colon and differed in bacterial biomass and microbiome structure.

In the present study, we compared the lethality of ICs in the mouse cecum and colon. The observations showed that cecal ICs produced a shorter median survival time, indicating that the cecum perforation might exhibit higher sepsis-associated lethality than the colon perforation. Moreover, we noticed that the blood biochemical indicators for liver impairment (ALT and AST), tissue damage (LDH), kidney dysfunction (BUN and Creatinine), and heart dysfunction (CK) were significantly higher or tended to increase in the FIP model developed with ICs from cecum which further corroborated by the histological examinations on liver, lung and kidney tissues. Compared to the colon group, we noticed more severe pathological alterations (including liver, kidney, and lung) in the cecum specimens than in the colon specimens, further corroborating the greater sepsis-associated lethality of cecum FIP mice. These findings are lined with clinical observations that the abdominal infection caused by intestinal perforation on the right side of the colon is more likely to have a poor prognosis and poor antibiotic treatment effect than on the left side [33] and a higher mortality rate (30-72%) occurred in cecum perforation [34]. In our study, the mice did not receive any treatment during sepsis progression, which differs from the clinical patients who were well treated with antibiotic administration. In short, the present study adopted a very objective strategy that did not introduce therapeutic interference to observe sepsis progression caused by different ICs of intestine segments. We assumed the cecum perforation led to more serious infection and severe inflammation than the colon perforation.

Further study on the blood plasma samples showed that the characteristic cytokine markers of sepsis[35], including TNFα, IL-1α, IL-1β, IL-6, and IL-10, increased significantly during the experimental timeline in the cecum-FIP mice compared to the colon group or the control mice. Similarly, we also demonstrate that levels of Eotaxin, G-CSF, GM-CSF, IFN-γ, IL-2, IL-3, IL-4, IL-5, IL-6, IL-9, IL-12(p40), IL-12(p70), IL-13, IL-17A, KC, MCP-1, MIP-1α, MIP-1β, RANTES rise significantly during cecum-induced sepsis but not significantly or less extent increase in colon-induced inflammation. These changes are often associated with inflammation in various organs, which indirectly indicates that cecum-induced sepsis has a higher systemic inflammatory response. IL-5 expression level is often associated with lung inflammation [36], and in the present study, we found plasma IL-5 raised in the Cecum-FIP mice that manifested more severe pneumonia and pathological changes. IL-6 is a sensitive marker of inflammation and can be of prognostic value in critically ill patients with multi-organ failure [37]. Interestingly, compared to colon-induced sepsis, more severe inflammation happened in cecum-induced sepsis. Patients with sepsis had a significant increase in circulating IFN-γ released by CD4^+^ T cells *via* IL-12 [38]. These increased plasma levels of injury and inflammatory markers confirmed the complexity of the dysregulated immune response during cecum-induced sepsis with exacerbated systematic inflammation and acute lung injury and worsened survival.

The composition of ICs varies in nutrients, pH, ions, gut microbes, and food debris in different segments, and their leakage into the peritoneal cavity can cause unexpected outcomes [1, 2, 27, 39]. The gut microbiome is the main component of the intestine and may vary in different sites [25, 26]. Moreover, the gut microbiome taxonomic differences are associated with additional mediators of disease pathogenesis, such as metabolites, short-chain fatty acids (SCFA), and metagenomic changes [1, 2, 39, 40]. Our study demonstrates that the cecum ICs harbor a ∼ 3-fold increased bacterial burden in unit weight than the colonic ICs, which could explain the increased illness severity and systemic inflammation due to increased infectious load and pathogen-associated molecular pattern (PAMPs) [2, 40]. In addition, we observed that the texture of the cecal contents is more fluid than that of the colon, which may indicate that more intestinal flora could quickly enter the abdominal cavity when the cecum is perforated. This high bacterial density and more fluidic characteristics in the cecum segment collectively lead to a more severe infection during cecal perforation. Although the alpha diversities of the cecum and colon microbiomes were similar in this study and suggested similar richness and diversity, these beta diversities demonstrated that the cecum and colon hold a different microbial community structure. Interestingly, several bacterial populations were identified in the microbial community as potential immunological or pathogenic [2]. For example, at the phylum level, we observed that the potentially pathogenic bacteria *p*_*Firmicutes* [41] primarily caused an increase in the cecum group.

We also noticed that several potentially pathogenic gut flora [42–44], including genera *Helicobacter*, *Mucispirillum, and Lachnospiraceae_FCS020_group*, and the family *Desulfovibrionaceae,* were more abundant in the cecum than in the colon group at the lower taxonomic level. Furthermore, inflammation-associated *Lachnoclostridium* and *Desulfovibrio* [45] also increased in the cecum compared to the colon group. In the present study, we knew *Helicobacter* could stimulate CD4^+^ T cells to Th1 differentiation through IL-12 production and enable these cells to secrete cytokines such as IL-1, IL-6, TNF-α and IFN-γ [46]; *Mucispirillum and Lachnospiraceae_FCS020_group* can promote intestinal inflammatory response [47, 48]; *Desulfovibrionaceae* were known for their sulfate-reducing capacities, as a result of this producing toxic hydrogen sulfide [49]. Therefore, these potentially pathogenic bacteria would explain their high IC-associated pathogenic potential for mouse survival and more severe multi-organ injury. In contrast, the potentially beneficial gut microbiome, such as the genera *Alloprevotella*, *Alistipes*, *Bacteroides*, *Parabacteroides*, *Rikenellaceae_RC9_gut_group*, *Prevotellaceae_UCG-001*, Prevotellaceae_NK3B31_group, and family *Muribaculaceae* [43, 44] decreased in the cecum group. In addition, we found that the *Firmicutes* to *Bacteroidota* (F/B) ratio [50–52] was higher in the cecum than in the colon group, indicating a dysbiosis in the community structure. Therefore, our results demonstrate that the greater accumulation of potentially pathogenic and less beneficial bacteria in the cecum has pro-inflammatory properties, causing more serious inflammatory storms and contributing to their lethality during the perforation or puncture.

In recent years, many updated international consensus definitions for sepsis and septic shock provide microbiota-targeted tools to improve sepsis-associated outcomes [53]. The target of changing antibiotics is to provide moderate homeostasis of the healthy intestinal microbiota. Therefore, we conducted a series of experiments starting with the different intestinal flora and demonstrated that causing abdominal infections was related to the ratio of different intestinal flora rather than just considering the presence of pathogenic bacteria. In the study, we compared the different IC-associated lethality for the first time while mimicking different perforation sites. The study showed that the cecal ICs are more lethal to mouse survival and cause multi-organ damage, which provides an understanding for evaluating and modulating the microbiota community to decrease the levels of harmful bacteria and increase the beneficial bacteria, thus minimizing antibiotic overuse. However, we did not compare the ICs of the cecum and colon in humans, and such clinical associations require further investigation. Of note, the perforations in the patients who received antibiotic administration make it challenging to compare to the animal data (without antibiotic treatment) in our study.

Our work suggested that the severity of cecum ICs-developed sepsis was associated with the bacterial burden and enriched pathogenic bacteria in microbial communities. However, we did not detect additional mediators, such as SCFA, fungal or other pathogens, and metagenomic changes, which need further investigation. Further work is needed to identify the pathogenic bacteria that drive sepsis and mechanistic study.

In summary, we standardized and developed a protocol to examine the different sites (cecum and colon) of perforation driving the sepsis. Our study demonstrated that cecum ICs developed more lethality, systemic inflammation, and multi-organ damage than colon ICs and found that the sepsis severity developed by perforation was associated with bacterial burden and increased abundance of potentially pathogenic bacteria in the cecum. This study provides the first experimental evidence to support that different perforations in the low digestive tract vary in terms of their prognoses. Therefore, it will be beneficial to design microbiota-based strategies to intervene in the microbial community to treat or slow the progression of sepsis or abdominal infections.

## 4. Materials and methods

### 4.1 Study design

To explore the possible cause of the severity of sepsis from different punctures or perforations in the lower digestive tract in mice, we used the FIP model [29] to evaluate gut microbes from the different intestinal sites and test whether they produced different effects on systematic inflammatory responses (SIRS) or survival of the mice. As shown in **Figure 1A**, the ICs from the cecum and colon (including the ascending, transverse, and descendant colon sections) were collected individually from the site of the euthanized mice (n = 10). Briefly, their abdominal were opened to expose different intestine fragments, and the ICs were scrutinized using sterile forceps to the clean tube. A small proportion of each IC sample was stored at −80 _ for subsequent 16S rRNA gene sequencing analysis. The remaining IC samples were mixed, weighed, and dissolved to form the solution for intraperitoneal injections into C57BL/6J mice for 16S rDNA quantification, survival, biochemical, cytokines and histological analysis.

### 4.2 Mice and fecal-induced peritonitis (FIP) model

The male C57/BL6J (7–9-week-old) mice were purchased from the Shanghai Model Organism Center (SMOC) and maintained for ∼2 weeks under a 12-h day/night specific-pathogen-free hood with free access to food and water for animal experiments. All procedures were carried out under the regulations for animal experimentation and were approved by the ethics committee of Tongren Hospital, Shanghai Jiao Tong University School of Medicine (No. A2022-012-01).

To assess the lethality and prognosis of different ICs, we adopted the FIP sepsis model [29] to construct the sepsis model in this study. Briefly, ICs from the different fragments of the cecum or colon were collected, followed by sacrifice and dissection of the donor mice (n = 10). ICs from 10 mice were mixed, weighed, and dissolved in saline at a 90 mg/ml concentration. Finally, the IC solutions were filtered with a 70-µm cell strainer and intraperitoneally injected (i.p.) with a dose of 2 g/kg body weight as previously described [29]. Then, we monitored the mouse survival and recorded their death time for a survival curve analysis among the saline (n=8), cecum (n=9), and colon (n=10) groups. They were euthanized at 28 h-post injection when the humane endpoints reached according to the following criteria: unable to crawl, unresponsive to touch stimulation, muscle trembling, arching hair on the back, dyspnea, and large amount of secretions in the mouth and nose. For histology analyses and biochemical analysis, mice (n=4 per group) were peritoneally injected with saline, cecum-ICs and colon-ICs to euthanization at 16h post-injections for dissecting tissues for further analysis as described below.

### 4.3 Tissue preparations, hematoxylin-eosin staining and pathological scoring

After intraperitoneally injected with the indicated ICs or saline for 16 h, the mice received anesthesia with 0.3% pentobarbital sodium followed by whole blood collection. Then, they were killed for tissue preparations. Freshly dissected livers, lungs, and kidneys were fixed in 4% paraformaldehyde to prepare paraffin-embedded sections. The 3 µm sections were used to perform the hematoxylin-eosin (H&E) staining as described previously [54], scanned by the Digital pathology slide scanners (KFBIO, Ningbo, China), and viewed by its K-ViEWER software. The clinical scores of the liver [55, 56], kidney [57], and lung [58] were employed by the clinic pathologist to evaluate the specific manifestations of pathological tissues.

### 4.4 Multiplex-Immunofluorescence staining and quantification

TSAPLus Fluorescence Triple Staining Kit (Wuhan Servicebio Technology Co., Ltd., Wuhan, China) was used for multiple immunofluorescence staining of lung specimens paraffin sections, i.e., herein sequentially labeling antigen with TNF-α (#GB11188, 1L1000), Ly6G (#GB11229, 1:2000) and ZO-1 (#GB111402, 1:100) antibodies purchased from Wuhan Servicebio Technology Co., Ltd. to evaluate the degree of lung injury when receiving the FIP. The slides were counterstained with 4’-6-diamidino-2-phenylindole (DAPI) and mounted with anti-fading solutions. The slides were scanned with the 3DHISTECH scanner and Viewer (Pannoramic MIDI, Hungary). The Ly6G- and TNFα− positive numbers and ZO-1 staining area were quantified using HighPlex FL v3.1. module or Area Quantification FL v2.1.2 module of the Indica Labs HALO software (v3.0.311.314) and plotted as the ratio to the Saline group.

### 4.5 Blood Biochemical Indicators Detection

The −80 °C stored plasma was detected for alanine aminotransferase (ALT), aspartate aminotransferase (AST), Creatinine, creatinine kinase (CK), Blood Urea Nitrogen (BUN), and lactate dehydrogenase (LDH) in the saline- and gut-microbes receiver group (n=3-4 per group). All biochemical indicators were measured using the Nanjing Jiancheng Bioengineering Institute (Nanjing, China) kit.

### 4.6 Cytokines measurements

The mouse plasma cytokines from the saline- and gut-microbes receiver mice (n=4 per group) were measured by Luminex liquid-phase suspension chip (Bio-Plex Pro Mouse Cytokine Grp I Panel 23-plex, #M60009RDPD, Wayen Biotechnologies (Shanghai), Inc). Briefly, the plasma and standard samples were diluted with Assay buffer to load on the chip for 30 min incubation. The chip incubates with the detection antibody for another 30 min, followed by Streptavidin-PE for 10 min. Finally, the multi-plate was detected by a Bio-Plex 200 system (Luminex Corporation, Austin, TX, USA) and the cytokine concentration was determined according to the standard curve fit.

### 4.7 RNA extraction, reverse transcription and quantitative PCR (qPCR)

The frozen kidney was homogenized for RNA extraction with Trizol Reagent (Invitrogen), quantification, reverse transcription, and SYBR green quantitative PCR for detecting the gene expression as previously [59–61]. The primers for qRT-PCR are listed below: *neutrophil gelatinase-associated lipocalin* (*NGAL*) (Forward: 5’-CTCAGAACTTGATCCCTGCC-3’, Reverse: 5’-TCCTTGAGGCCCAGAGACTT-3’); *Actb* (Forward: 5’-GGCTGTATTCCCCTCCATCG-3’, Reverse: 5’-CCAGTTGGTAACAATGCCATGT-3’). The mRNA expression was normalized to the *Actb* mRNA and interpreted as the foldchange of the Control group (Saline).

### 4.8 16S rDNA qPCR

16S rDNA from mouse cecum ICs and colonic ICs was analyzed by SYBR green (#Q711-03, Vazyme, Nanjing, China) qPCR for bacterial 16S rDNA genes using universal primers 341F (5’-cctacgggnggcwgcag-3’) and 785R (5’-gactachvgggtatctaatcc-3’) according to the following cycling conditions [62]: 50°C for 2 min, 95°C for 2 min, 45 cycles of 95°C for 15 s, 58°C for 15 s and 72°C for 1 min. Quantitation of 16S rDNA copies normalized to pUC57 vector harboring 16S V3 and V4 region and interpreted as the copies per fecal (in mg).

### 4.9 16S rRNA gene sequencing analysis

We performed 16S rRNA gene sequencing analysis as described previously [63]. Briefly, IC samples from the cecum or colon fragments of the donors (9–11-week-old male mice) were collected and stored at −80 °C after collection. According to the manufacturer’s instructions, bacterial DNA was isolated from the IC samples using a MagPure Soil DNA LQ Kit (Magen, Guangdong, China), measured using a NanoDrop 2000 spectrophotometer (Thermo Fisher Scientific, Waltham, MA, USA), and evaluated by electrophoresis. First, the bacterial 16S rRNA gene was amplified using universal primer pairs that targeted the V3-V4 hypervariable regions (343F: 5′-TACGGRAGGCAGCAG-3′; 798R: 5′-AGGGTATCTAATCCT-3′). The amplification products were then purified with Agencourt AMPure XP beads (Beckman Coulter Co., USA) and quantified using a Qubit dsDNA assay kit. Finally, the DNA was sequenced using the Illumina NovaSeq6000 platform (Illumina Inc., San Diego, CA) by OE Biotech Company (Shanghai, China).

Paired-end raw reads were deposited in the NCBI Sequence Read Archive (SRA) with BioProject ID accession number PRJNA872016, preprocessed using Trimmomatic software [64], and assembled using FLASH software [65]. Reads with 75% of bases above Q20 were retained using QIIME software (version 1.8.0) [66]. The clean reads were then subject to primer sequence removal and clustering to generate operational taxonomic units (OTUs) using VSEARCH software with a 97% similarity cutoff [67]. The representative read of each OTU was selected using the QIIME package. All representative reads were annotated and blasted against the Silva database (Version 132) using the RDP classifier (confidence threshold was 70%) [68]. The alpha diversity, including the Chao1 index [69], Shannon index [70], phylogenetic diversity (PD) tree, and beta diversity principle coordinate analysis (PCoA) with a Bray–Curtis distance matrix, were determined using the QIIME software. Linear discriminant analysis effect size (LEfSe) analysis was performed to identify unique microbial composition species associated with infection severity. Microbiota with a linear discriminant analysis (LDA) score greater than 3.5 were defined as different genera.

### 4.10 Statistical Analysis

Statistical analysis was performed on raw data for each group using a One-way ANOVA with post unpaired t-test, Kruskal-Wallis test, or Mantel–Cox test as indicated using GraphPad Prism 8 software (GraphPad Software, San Diego, CA). Data were expressed as mean ± standard error of the mean (SEM). P-values refer to the probability of the null hypothesis that the means do not differ. A p < 0.05 was considered significant, and p ≥ 0.05 was not significant.

## Ethics approval

The animal study (No. A2022-012-01) was reviewed and approved by the ethics committee of Tongren Hospital, Shanghai Jiao Tong University School of Medicine.

## Consent for publication

All authors agree to publication.

## Availability of data and materials

The raw reads of 16S rRNA-seq have been deposited in the SRA database and assigned a BioProject accession number PRJNA872016 for 16S rRNA-seq. In addition, other raw data and materials can be reasonably requested from the authors (<u>Y. Lu or J. Zhang</u>).

## Authors’ contributions

Y. Lu, J. Zhang, and K. Xu designed the study. K. Xu, J. Tan, D. Lin and Y. Lu performed experiments. K. Xu, J. Tang, D. Lin, Y. Chu, L. Zhou, J. Zhang and Y. Lu analyzed the data. Y. Lu and J. Zhang supervised the study. Y. Lu, K. Xu, J. Tan, Y. Chu, L. Zhou, and J. Zhang wrote the paper. All authors reviewed the results and approved the final version of the manuscript.

## Acknowledgements

We thank OE Biotech Co., Ltd. (Shanghai, China) for assistance in the 16S RNA gene sequencing analysis and their cloud tools for data analysis.

## Supplementary information

The Supplementary information for this article can be found online.

