## Supplementary material for "Gut microbes of the cecum versus the colon drive more severe lethality and multi-organ damage": Supplemantal Figure 1-2 and Table 1-2: Supplementary information_BIORXIV.docx

**Supplementary Figure S1**

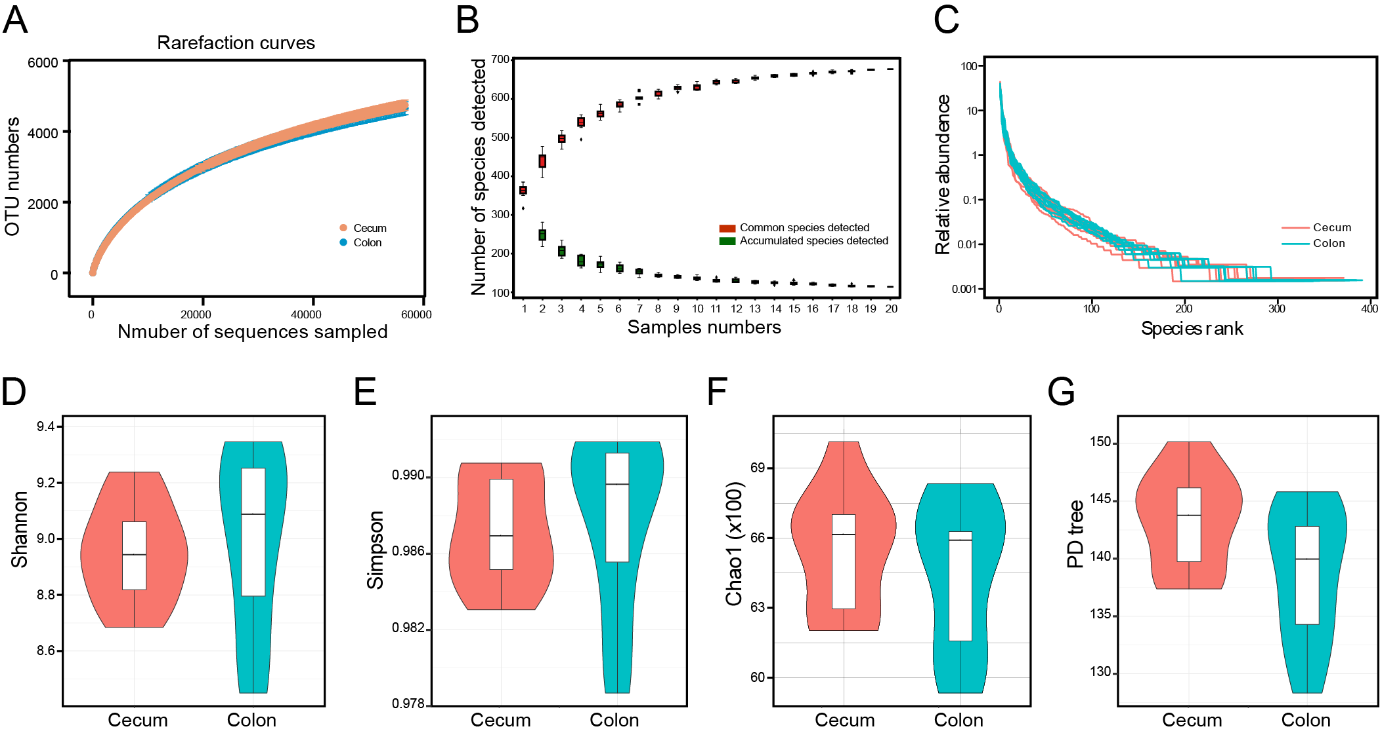
**Supplementary Figure S1.** (**A**) Rarefaction curves based on OTUs of cecum and colon. (**B**) The numbers of species detected curve. (**C**) Relative abundance of species rank curve. (**D-G**) The alpha-diversity index is shown by Shannon (**D**), Simpson (**E**), Chao1 (**F**), and the phylogenetic diversity (PD) tree (**G**).

**Supplementary Figure S2**

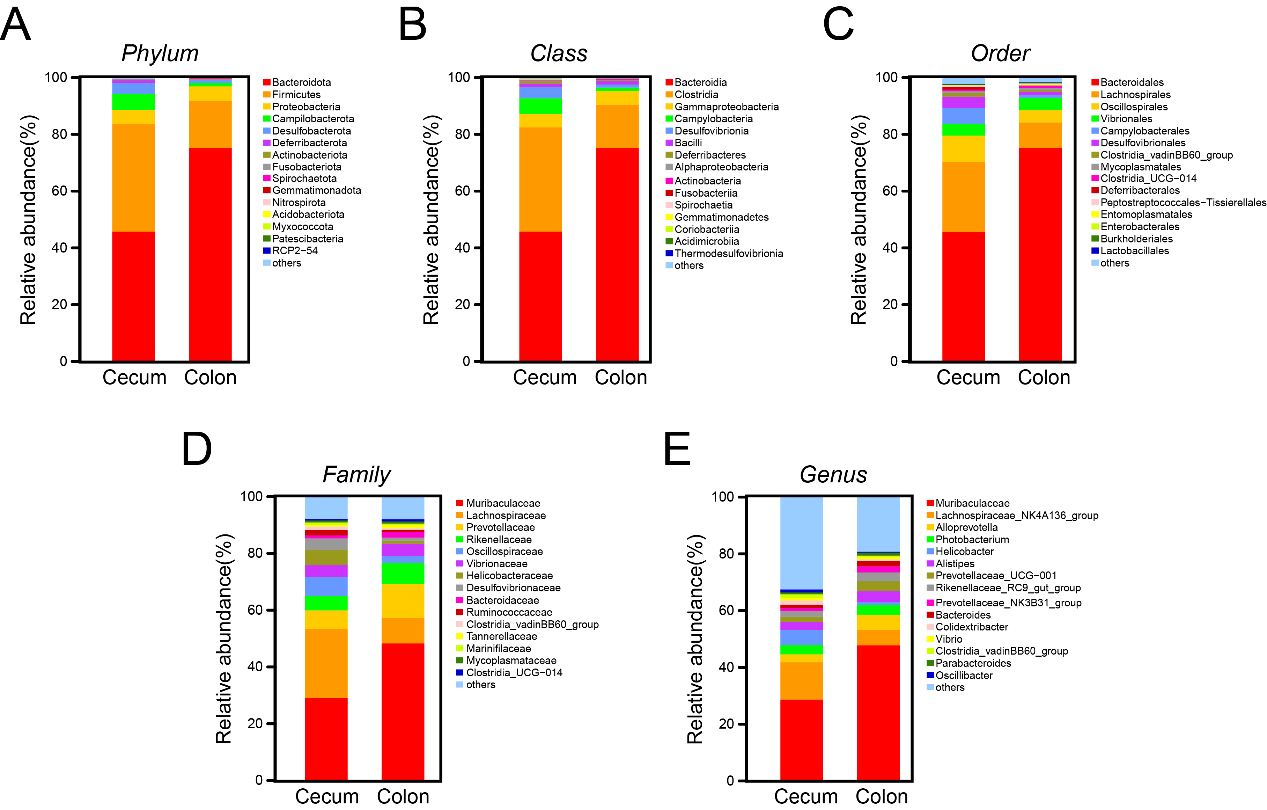
**Supplementary Figure S2.** Relative abundance of microbiome community based on *phylum* (**A**), *class* (**B**), *order* (**C**), *family* (**D**), and *genus* (**E**).

**Supplementary Table S1**. Summary of the tags and OTU after taxonomic assignment

| Sample_ID | clean_tags | valid_tags | valid_percent | OTU_counts | Total_OTUs |
| --- | --- | --- | --- | --- | --- |
| Cecum_1 | 72496 | 59292 | 81.79% | 4884 | 17172 |
| Cecum_2 | 75921 | 64715 | 85.24% | 5068 | 17172 |
| Cecum_3 | 77937 | 66962 | 85.92% | 4762 | 17172 |
| Cecum_4 | 74613 | 63174 | 84.67% | 5084 | 17172 |
| Cecum_5 | 77056 | 65523 | 85.03% | 4993 | 17172 |
| Cecum_6 | 77356 | 66192 | 85.57% | 4725 | 17172 |
| Cecum_7 | 75612 | 63878 | 84.48% | 5136 | 17172 |
| Cecum_8 | 76747 | 62609 | 81.58% | 4648 | 17172 |
| Cecum_9 | 72964 | 56755 | 77.78% | 4884 | 17172 |
| Cecum_10 | 76838 | 63846 | 83.09% | 5229 | 17172 |
| Colon_1 | 73432 | 63362 | 86.29% | 5014 | 17172 |
| Colon_2 | 73989 | 65241 | 88.18% | 4865 | 17172 |
| Colon_3 | 75034 | 65383 | 87.14% | 4525 | 17172 |
| Colon_4 | 73811 | 63566 | 86.12% | 5145 | 17172 |
| Colon_5 | 75327 | 65138 | 86.47% | 4764 | 17172 |
| Colon_6 | 74092 | 63782 | 86.08% | 4476 | 17172 |
| Colon_7 | 76033 | 65680 | 86.38% | 5164 | 17172 |
| Colon_8 | 76260 | 67299 | 88.25% | 5157 | 17172 |
| Colon_9 | 75089 | 67929 | 90.46% | 4728 | 17172 |
| Colon_10 | 75682 | 66623 | 88.03% | 5192 | 17172 |

**Supplementary Table S2 Summary of Adonis statistical analysis**

| Adonis | F.Model | R2 | p-value(>F) | Signif |
| --- | --- | --- | --- | --- |
| adonis.binary_jaccard | 2.0074 | 0.10033 | 0.001 | *** |
| adonis.bray_curtis | 4.8997 | 0.21396 | 0.001 | *** |
| adonis.euclidean | 3.3166 | 0.15559 | 0.001 | *** |
| adonis.unweighted_unifrac | 2.1336 | 0.10597 | 0.001 | *** |
| adonis.weighted_unifrac | 24.817 | 0.57961 | 0.001 | *** |
